# A new gene encoding a cytosolic glutamine synthetase in pine is linked to developing tissues

**DOI:** 10.1101/2022.10.27.514046

**Authors:** José Miguel Valderrama-Martín, Francisco Ortigosa, Juan Carlos Aledo, Concepción Ávila, Francisco M. Cánovas, Rafael A. Cañas

## Abstract

The enzyme glutamine synthetase (EC 6.3.1.2) is mainly responsible for the incorporation of inorganic nitrogen into organic molecules in plants. In the present work, a new pine *GS1* (*PpGS1b.2*) gene was identified, showing a high sequence identity with the *GS1b.1* gene previously characterized in conifers. Phylogenetic analysis revealed that the presence of *PpGS1b.2* is restricted to the genera *Pinus* and *Picea* and is not found in other conifers. Gene expression data suggest a putative role of *PpGS1b.2* in plant development, similar to other *GS1b* genes from angiosperms, suggesting evolutionary convergence. The characterization of GS1b.1 and GS1b.2 at the structural, physicochemical, and kinetic levels has shown differences even though they have high sequence homology. Alterations in the kinetic characteristics produced by the site-directed mutagenesis approach carried out in this work strongly suggest an implication of amino acids at positions 264 and 267 in the active center of pine GS1b.1 and GS1b.2. Therefore, the amino acid differences between GS1b.1 and GS1b.2 would support the functioning of both enzymes to meet distinct plant needs.

## INTRODUCTION

Nitrogen (N) is an essential element, a constituent of the main biomolecules and a limiting factor for plant growth (Hirel and Krapp, 2021). N is assimilated from ammonium into organic molecules by the glutamine synthetase (GS, EC 6.3.1.2)/glutamate synthase (GOGAT, EC 1.4.7.1) cycle. Ammonium is first incorporated into glutamate to form glutamine in an ATP-dependent reaction catalyzed by the GS enzyme (Heldt and Piechulla, 2011), and then this glutamine together with 2-oxoglutarate is used to produce two glutamate molecules by the GOGAT enzyme (Bernard and Habash, 2009). Studies have shown that up to 95% of ammonium is assimilated via the GS/GOGAT cycle (Lea *et al*., 1999) for the formation of glutamine and glutamate, which, in turn, will be used to produce all N-containing biomolecules in the plant (Forde and Lea, 2007; Bernard and Habash, 2009).

The GS enzyme has been widely studied in plants since it is directly responsible for the incorporation of inorganic N into organic molecules. Recently, three different lineages of *GS* genes have been identified in seed plants: *GS1a* and *GS1b* encode cytosolic enzymes, and *GS2* encodes a plastid-located enzyme (Valderrama-Martín *et al*., 2022). The three *GS* gene lineages are present in cycads and *Ginkgo biloba*, as well as basal angiosperms. Nevertheless, no GS2 genes have been found in other gymnosperms, such as conifers and gnetales, and no *GS1a* genes have been found in modern angiosperms, including monocot and eudicotyledon species (Valderrama-Martín *et al*., 2022). In general, GS1b is encoded by a small multigene family, while GS1a and GS2 are usually encoded by a single nuclear gene (James *et al*., 2018; Valderrama-Martín *et al*., 2022).

GS2 and GS1a are associated with photosynthetic organs (Blackwell *et al.*, 1987; Ávila *et al.*, 2001), and their expression is regulated by light conditions (Cantón *et al.*, 1999; Gómez-Maldonado *et al.*, 2004a; Valderrama-Martín *et al.*, 2022). Indeed, GS2 and GS1a are considered to play a fundamental role in the assimilation of the ammonium released during photorespiration and nitrate photoassimilation processes (Wallsgrove *et al.*, 1987; Blackwell *et al.*, 1987; Cantón *et al.*, 1999; Tegeder and Masclaux-Daubresse, 2017). In this sense, new evidence suggests that the *GS2* gene may arose through a gene duplication from a *GS1a* gene in a common ancestor of cycads, ginkgo, and angiosperms (Valderrama-Martín *et al*., 2022).

GS1b corresponds to the GS1 isoenzyme traditionally studied in model angiosperms. Although this lineage is represented by a unique gene in most of the gymnosperms, in ginkgo and angiosperms, GS1b is represented by a small multigenic family. These genes have different expression patterns depending on the organ and physiological conditions accounting for their different functions (Hirel and Krapp, 2021). These enzymes have been described as a key components of plant nitrogen use efficiency, with essential roles in processes such as senescence (Thomsen *et al*., 2014), amino acid catabolism, primary assimilation, and different stress responses (Bernard and Habash, 2009). The different genes of this lineage are differentially regulated by developmental state, tissue, nutritional status, and external stimuli (Thomsen et al. 2014; Hirel and Krapp, 2021). Finally, several studies have focused on the enzymatic characterization of GS from angiosperms and gymnosperms (Sakakibara *et al*., 1996; de la Torre *et al*., 2002; Ishiyama *et al*., 2004a; Ishiyama *et al*., 2004b; Ishiyama *et al*., 2006; Yadav, 2009; Zhao *et al*., 2014; Castro-Rodríguez *et al*., 2015) to define a more accurate role landscape for the different GS isoforms.

Some GS1b isoforms are directly related to developmental processes and have been associated with grain yield in crops. *AtGS1.1* and *AtGS1.2* from *Arabidopsis thaliana* are involved in seed production and germination (Guan *et al*., 2015). *AtGS1.1* has also been described to be involved in root development during seed germination and *AtGS1.2* plays a role in rosette development (Lothier *et al*., 2011; Guan *et al*., 2015). Indeed, a recent study over of *AtGS1.1*, *AtGS1.2* and *AtGS1.3 Arabidopsis* mutants suggested synergistic roles for these genes in plant growth and development (Ji *et al*., 2019). In cereals, enzymes of this GS lineage are involved in seed yield and plant development, such as *GS1;3* from *Oryza sativa* and *Hordeum vulgare*, which play roles in seed maturation and germination (Goodall *et al*., 2013; Fujita *et al*., 2022). Thus, overexpressing lines of *HvGS1.1* showed an improvement in grain yield (Gao *et al*., 2019). Rice mutants lacking the *OsGS1;1* gene presented reduced grain filling and growth (Tabuchi *et al*., 2005), although the same phenotype was present in rice lines overexpressing *OsGS1;1* (Bao *et al*., 2014). In addition, rice lines grown in culture chambers and overexpressing *OsGS1;1* presented an increase in spikelet yield. Rice mutants for *OsGS1b;2* also presented a depletion in the number of tillers (Funayama *et al*., 2013), and *Sorghum bicolor* lines overexpressing *GS1* genes exhibited the opposite phenotype (Urriola and Rathore, 2015). Studies in *Zea mays* using mutant lines for *ZmGS1b.3* and *ZmGS1b.*4 have shown the roles of these genes in kernel number and size, respectively (Martin *et al*., 2006). Transgenic lines of *Phaseolus vulgaris* overexpressing GS1 also showed earlier flower and seed development, while overexpressing GS1 lines of wheat showed an increase in grain weight (Habash *et al*., 2001). Moreover, a recent study on wheat indicated that *TaGS1.1* and *TaGS1.3* are mainly expressed in embryos and grain transport tissues, where these isoforms synergistically carry out ammonium assimilation (Wei *et al*., 2021).

In conifers, only one isoform of the GS1b family has been identified to date. The unique GS1b identified in conifers has been suggested to play an essential role in N remobilization to developing organs (Suárez *et al*., 2002). Previous works in pine have shown that GS1b is involved in the canalization of ammonium into glutamine during seed germination and the early developmental stages of seedlings (Ávila *et al*., 2001), which could be important for the loss of seed dormancy (Schneider and Gifford, 1994). Indeed, the roles of GS1b in seed development and germination are also supported by its expression patterns associated with the vascular system of zygotic and somatic pine embryos at different developmental stages and by its expression in procambium cells of pine zygotic embryos (Pérez-Rodríguez *et al*., 2005). Moreover, the expression of this isoenzyme has been suggested to be controlled by gibberellic acid, a phytohormone involved in many aspects of plant growth and development (Gómez-Maldonado *et al*., 2004b).

In this work, a new gene encoding a cytosolic GS (*PpGS1b.2*) was identified in maritime pine (*Pinus pinaster*). This gene was discovered through sequence searches in transcriptomic data from isolated tissues through laser capture microdissection (Cañas *et al*., 2017). Orthologs of this gene have also been identified in the genomes of other conifers, and phylogenetic analysis has revealed that *PpGS1b.2* belongs to the *GS1b* lineage. Although this new *GS1* gene presents a high sequence homology to the already known *PpGS1b*, hereafter *PpGS1b.1*, *PpGS1b.2* showed low expression levels with characteristic and localized tissue expression. The expression patterns suggest that this new gene could play a specific role during plant development, mainly during embryo development, as has been shown for other *GS1b* genes in angiosperms. Furthermore, a detailed comparative analysis of the kinetic properties of the isoenzymes GS1b.1 and GS1b.2 and single/double-point mutants of both isoforms support distinct functions for these enzymes in pine.

## RESULTS

### Sequence and phylogenetic analyses

A new cytosolic *GS* gene was identified in a transcriptomic analysis of tissues isolated using laser capture microdissection (Cañas *et al*., 2017). At the amino acid sequence level, the new GS presents 80.85% and 92.68% identity with PpGS1a and PpGS1b, respectively (Figure 1A). Despite the high identity between the coding sequences of the new gene and *PpGS1b*, the promoter regions of both genes are very distinct (Figure S1A). The lengths of the three pine GS proteins are very similar, with 357 residues for PpGS1a, 355 for PpGS1b and 357 for the new protein (Figure 1A). However, the calculated isoelectric points were more different between the pine GS proteins, being 6.21 in the case of PpGS1a, 5.73 for PpGS1b and 5.36 for the new protein.

**Figure 1.**
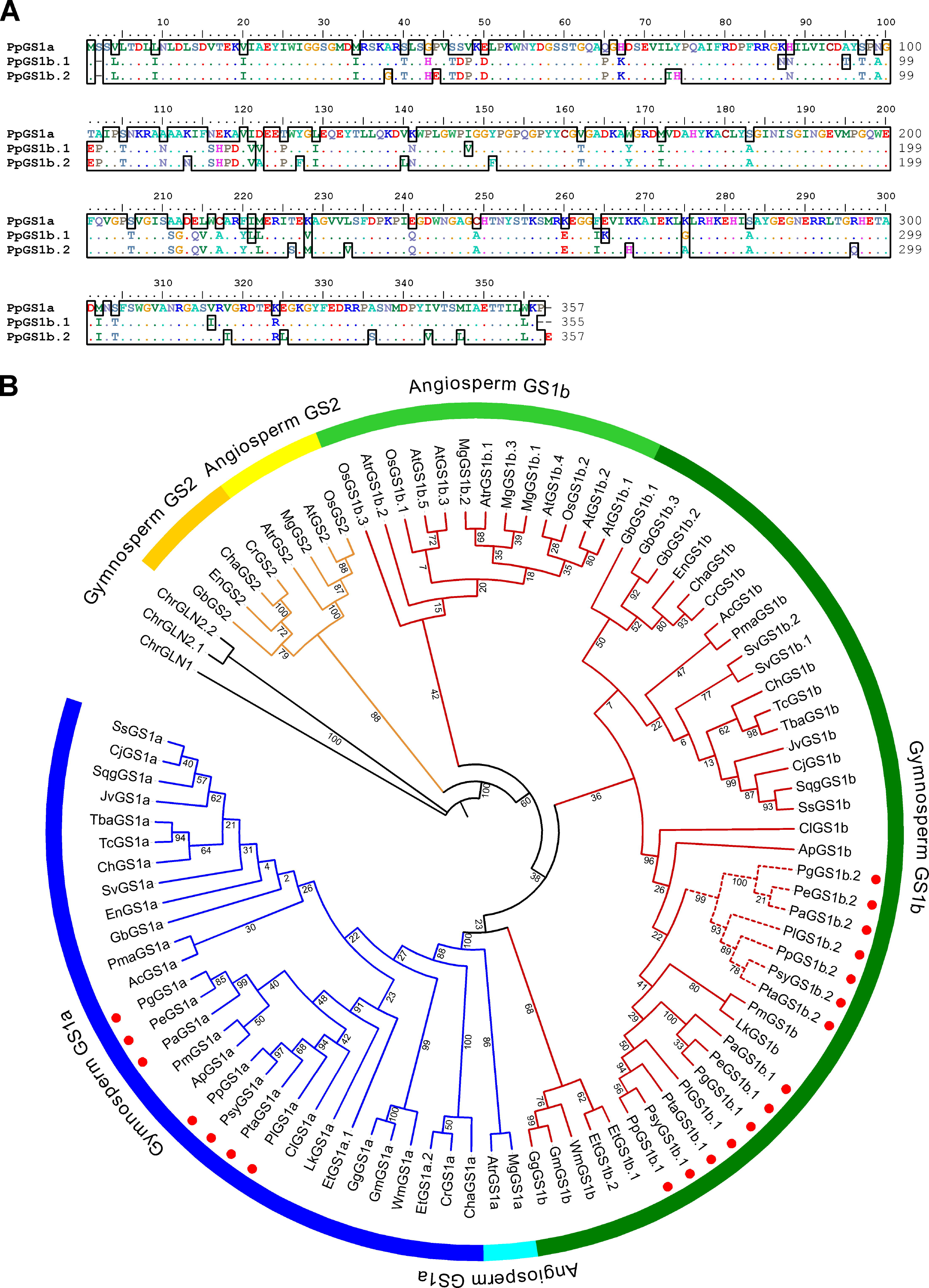
Protein alignment and evolutionary analysis by Maximum Likelihood method. **A.** Protein alignment of maritime pine GSs. PpGS1a sequence is showed as reference, dots highlight conserved residues in the three sequences. **B.** The evolutionary history was inferred by using the Maximum Likelihood method and JTT matrix-based model (Jones et al., 1992). The tree with the highest log likelihood (−12198.89) is shown. The percentage of trees in which the associated taxa clustered together is shown next to the branches. Initial tree(s) for the heuristic search were obtained automatically by applying Neighbor-Join and BioNJ algorithms to a matrix of pairwise distances estimated using the JTT model, and then selecting the topology with superior log likelihood value. The tree is drawn to scale, with branch lengths measured in the number of substitutions per site. This analysis involved 96 amino acid sequences. All positions containing gaps and missing data were eliminated (complete deletion option). There was a total of 348 positions in the final dataset. Evolutionary analyses were conducted in MEGA11 (Tamura et al., 2021). Numbers close to the branches shown bootstrap values. The first two letters of the sequence names correspond to the genera and species listed in Table □ S1. Golden tree branches correspond to GS2 sequences; blue branches to GS1a sequences; and red branches to GS1b sequences. Discontinuous lines in GS1b branches highlight the new sequences found in *Pinus* and *Picea* genera. Red dots shown the sequences from *Pinus* and *Picea* genera.

A phylogenetic analysis confirmed the classification of the GS from seed plants into three main groups, GS2, GS1a and GS1b, in line with previously reported results (Valderrama-Martín *et al*., 2022) (Figure 1B). As expected, no GS2 sequence was detected in conifers, but only those of GS1a and GS1b (Figure 1B). The new GS isoform was grouped within the conifer GS1b sequences; thus, the gene coding this new GS1b isoenzyme has been named *PpGS1b.2*. Orthologs of *PpGS1b.2* have also been detected in other members of the *Pinaceae* family of the genera *Pinus* and *Picea* but not in the rest of the conifers included in this analysis (Figure 1B).

### Gene expression analyses

The expression of *GS* genes in *P. pinaster* has been analyzed in different tissues and conditions to establish a framework that allows us to unravel the potential role of *PpGS1b.2* by comparing its expression pattern to other *GS* genes in maritime pine.

The expression profiles were analyzed in embryos and seedlings during the initial developmental stages (Figure 2A). *PpGS1a* expression was high in cotyledons and needles, lower in hypocotyls and nearly undetectable in roots and embryos except for germinated embryos. *PpGS1b.1* and *PpGS1b.2* expression patterns in embryos were very similar, with a peak of expression in germinated embryos. In seedlings, the expression was ubiquitous in all organs for both genes, although *PpGS1b.2* expression levels were lower than those of *PpGS1b.1*, between 5- and 10-fold. This expression pattern was different when isolated tissues were considered (Figure 2B). *PpGS1b.1* was expressed at high levels throughout the plant, especially in the root cortex, where the expression was 40 times that shown by this gene in the other samples. However, *PpGS1b.2* expression was very localized, mainly in the shoot apical meristem, emerging needles, developing root vascularization and root meristem. Expression was almost undetectable in the rest of the tissues analyzed. Finally, the expression of *PpGS1a* was detected only in the three photosynthetic tissues: the mesophyll of young needles, the mesophyll of cotyledons and the hypocotyl cortex.

**Figure 2.**
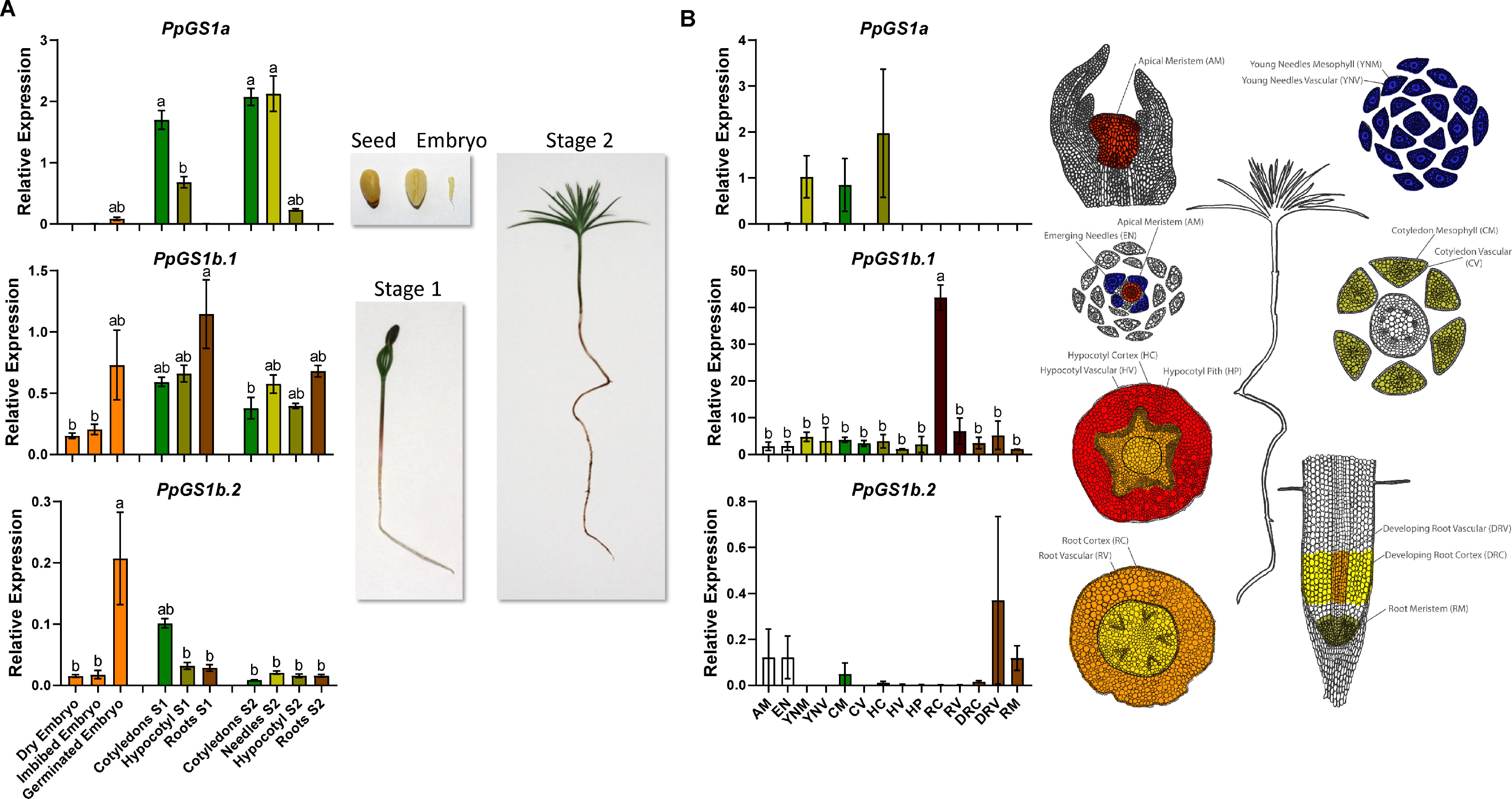
*GS* gene expression in maritime pine seedlings. **A.** Expression levels of *GS* genes of maritime pine during germination and initial seedling development. Stage 1 (S1) corresponds to seedlings with active mobilization of reserves from megagametophyte to the seedling (one-week-old from emergence). Stage 2 (S2) corresponds to seedlings without megagametophyte and developing the first new needles (one-month-old from emergence). **B.** Gene expression levels of GS in tissues from one-month-old seedlings (Cañas et al., 2017). AM, shoot Apical Meristem; EN, Emerging Needles; YNM, Young Needles Mesophyll tissue; YNV, Young Needles Vascular tissue; CM, Cotyledon Mesophyll tissue; CV, Cotyledon Vascular tissue; HC, Hypocotyl Cortex; HV, Hypocotyl Vascular tissue; HP, Hypocotyl Pith; RC, Root Cortex; RV, Root Vascular tissue; DRC, Root Developing Cortex; DRV, Root Developing Vascular tissue; RM, Root apical Meristem. Letters above the columns highlight the statistical significance (*P*<0.05) in a Tukey post-hoc test after an ANOVA analysis. Error bars show SE with n=3.

The seasonal expression of the three *GS* genes has also been quantified in needles from adult trees (Figure 3A). *PpGS1a* showed the highest expression, followed by *PpGS1b.1*, which was expressed between 10 and 30 times less than *PpGS1a*. The expression levels of *PpGS1b.2* were very low compared to those of other *GSs*. The expression patterns of the three genes in different whorls were as before, with higher levels in the first months of the year and lower levels at the end of the year. There was a remarkable exception for whorl 0 in May, the first harvesting month for the needles that emerged during the sampling year. *PpGS1b.2* exhibited an expression peak in whorl 0 in May. In contrast, *PpGS1a* had its lowest expression, and *PpGS1b.1* was expressed at similar levels to the other whorls. The relative abundance of *PpGS1b.2* transcripts was still one and two orders of magnitude lower than those of *PpGS1b.1* and *PpGS1a*, respectively. According to these results, the expression levels of the three genes were also analyzed in buds and emerging needles (Figure 3B-D). The expression of *PpGS1a* was almost undetectable in buds, but its expression rapidly increased in nascent needles by the end of the month. *PpGS1b.1* expression remained almost invariable in both organs with a similar expression pattern. The levels of *PpGS1b.2* were higher in the buds and decreased from Day 14 to 28 when the expression was similar in buds and emerging needles. The relative abundance of *PpGS1b.1* transcripts was still higher than that of *PpGS1b.2*.

**Figure 3.**
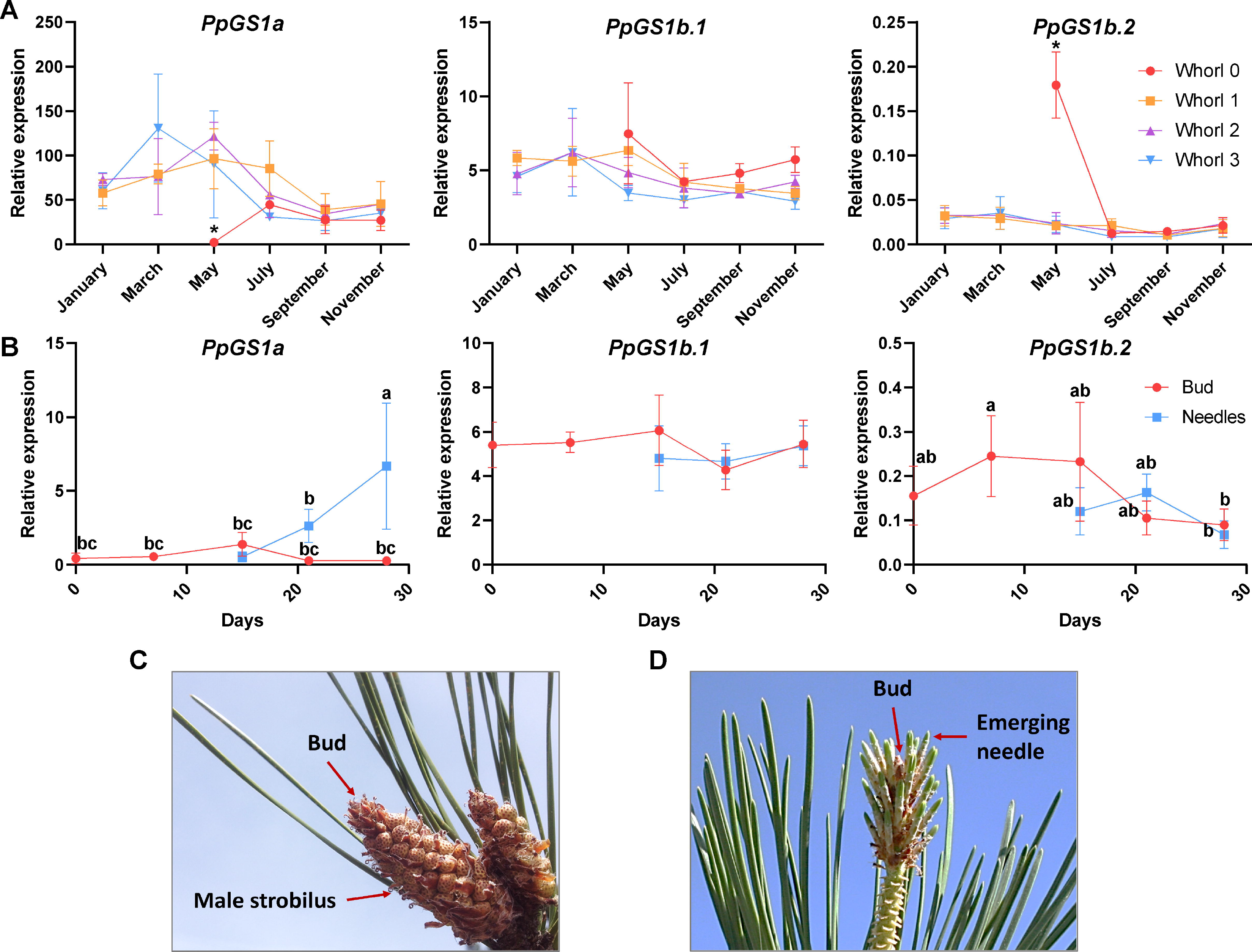
Seasonal *GS* expression profiles in pine needles from adult trees. **A.** Expression levels of *GS* genes were determined in needles from maritime pine along a year. Each needle whorl corresponds to the annual growth of a single year, the whorls were named by numbers, from 0 to 3 being this the oldest whorl. The whorl 0 corresponds to needles emerged in the same year of harvesting. For supplementary information see Cañas et al. (2015). Asterisks above the data points highlight the statistical significance (*P*<0.05) between needle whorls in a specific month in a Tukey post-hoc test after an ANOVA analysis. Error bars show SE with n=3. **B.** Expression levels of *GS* genes in buds and developing needles during the first 21 days of emergence. Letters above the data points highlight the statistical significance (*P*<0.05) in a Tukey post-hoc test after an ANOVA analysis. Error bars show SE with n=3. **C.** Picture of buds and male strobilus in April during the first harvesting time. **D.** Picture of buds and emerging needles in May at four harvesting point (21 days).

*GS* gene expression has also been analyzed at different developmental stages, including juvenile and mature xylem and phloem, as well as the male and female reproductive structures, different root zones and different stages of zygotic embryo development (Figure 4). In all those samples, *PpGS1a* expression was barely detectable. An example of *PpGS1a* expression is shown for phloem, xylem, and male and female strobili, with very low levels (< 0.04), even in female strobilus with an expression peak (< 0.08) (Figure 4A). *PpGS1b.1* expression was the highest observed thus far among the *GS* genes analyzed in vascular tissues and strobili (Figure 4A). Interestingly, *PpGS1b.2* expression was almost undetectable in vascular tissues, but its levels peaked in the male strobilus (approximately 0.28), opposite to what occurred with *PpGS1b.1* in that organ. In root samples, *PpGS1b.1* and *PpGS1b.2* presented a similar expression pattern, with increased expression in lateral roots and root tips, although the expression levels for *PpGS1b.1* were approximately 80-fold higher than that shown by *PpGS1b.2* (Fig, 4B). Finally, in zygotic embryos, the expression levels of both genes were significantly higher in the precotyledonary and early cotyledonary stages, where *PpGS1b.2 levels* were higher than those shown by *PpGS1b.1* (Figure 4C). However, this ratio of the expression of both genes was reversed in the later stages of development in cotyledonary and mature embryos. Nevertheless, the differences in expression between the two genes were not statistically significant in either case.

**Figure 4.**
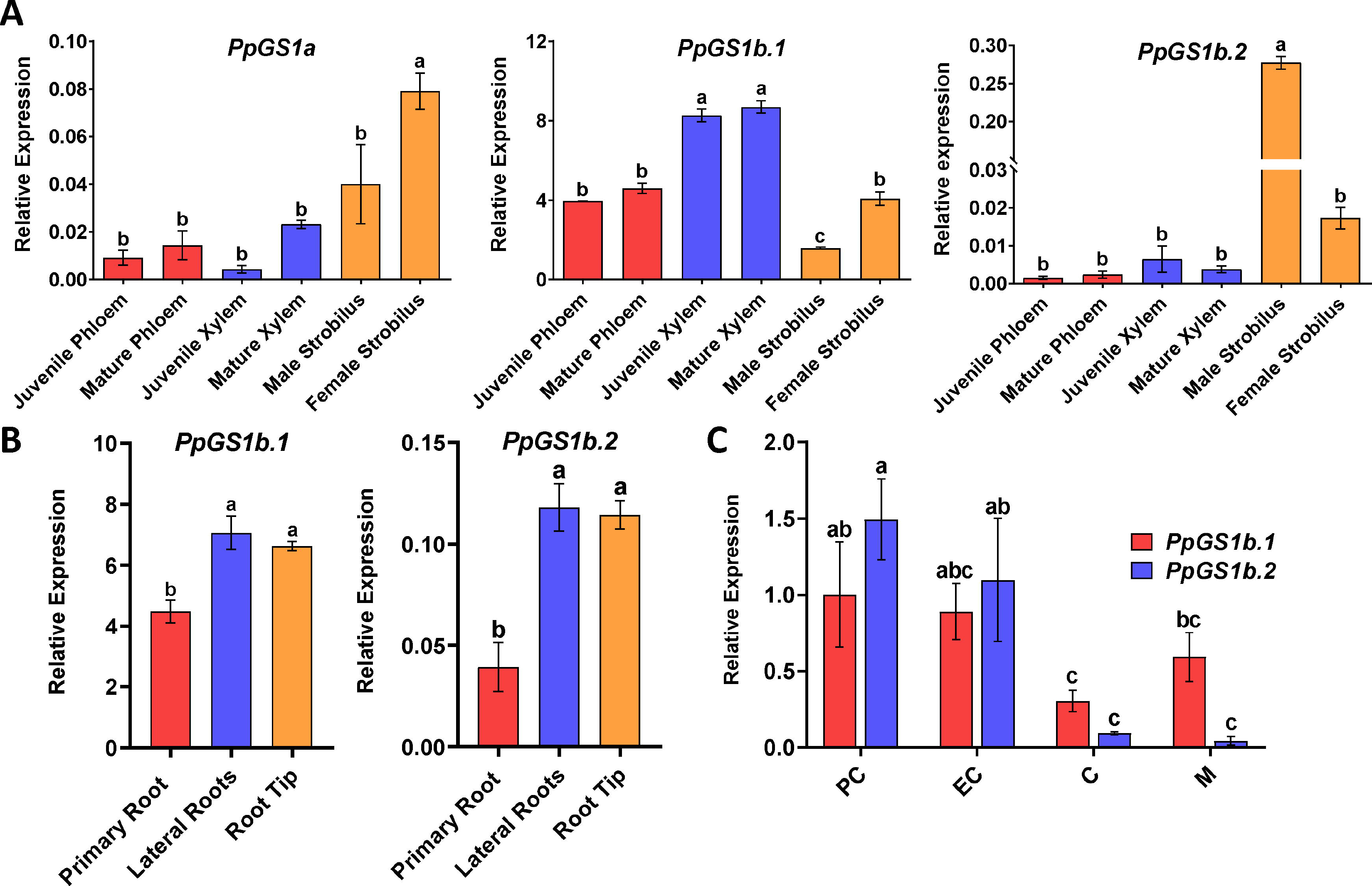
*GS* expression levels in different developing tissues. **A.** Gene expression levels of *PpGS1a*, *PpGS1b.1* and *PpGS1b.2* in different tissues of adult trees: juvenile and mature phloem; juvenile and mature xylem; and male and female strobili. **B.** Gene expression of *PpGS1b.1* and *PpGS1b.2* in different parts of the root from one-month-old seedlings: primary root, lateral roots, and root tip. **C.** Genes expression of *PpGS1b.1* and *PpGS1b.2* in different developmental stages of zygotic embryos: PC (pre-cotyledonary stage); EC (early cotyledonary stage), C (cotyledonary stage) and M (mature embryo). Letters above the columns highlight the statistical significance (*P*<0.05) in a Tukey post-hoc test after an ANOVA analysis. Error bars show SE with n=3.

### Protein structure prediction and physicochemical and kinetic properties

Very few differences were observed between the GS1b.1 and GS1b.2 subunit structures due to the similarity of their amino acid sequences (Figure5A,B). Both proteins presented a predicted decameric structure formed by two pentameric rings with small differences in structure and the disposition of the subunits in the quaternary structure (Figure S2A,B). However, the thermodynamic stability of GS1b.1 monomers was three times higher than that of GS1b.2 monomers (Table 1). The *in silico* replacement of residues of the GS1b.1 and GS1b.2 amino acid sequences displayed some differences in the structural stability of both enzymes (Figure S3). Some of the amino acids used for this analysis did not cause any notable effects on the structure or destabilized both proteins equally. However, several amino acids gave rise to large differences in the free energy of folding. Specifically, the inclusion of arginine or glutamate around position 280 produced a great destabilization of the structure of GS1b.2 but not of GS1b.1. Some of these amino acids also caused great destabilization of GS1b.2 when substituted at position 148 but did not have the same effect in GS1b.1. In fact, only isoleucine and arginine produced marked effects on the structural stability of GS1b.1. As small differences in the structure suggested that there might be changes in the physicochemical and kinetic properties of both enzymes, a functional comparison of the recombinant isoforms of GS1b.1 and GS1b.2 was performed (Figure 5, S4; Tables 1, 2).

**Figure 5.**
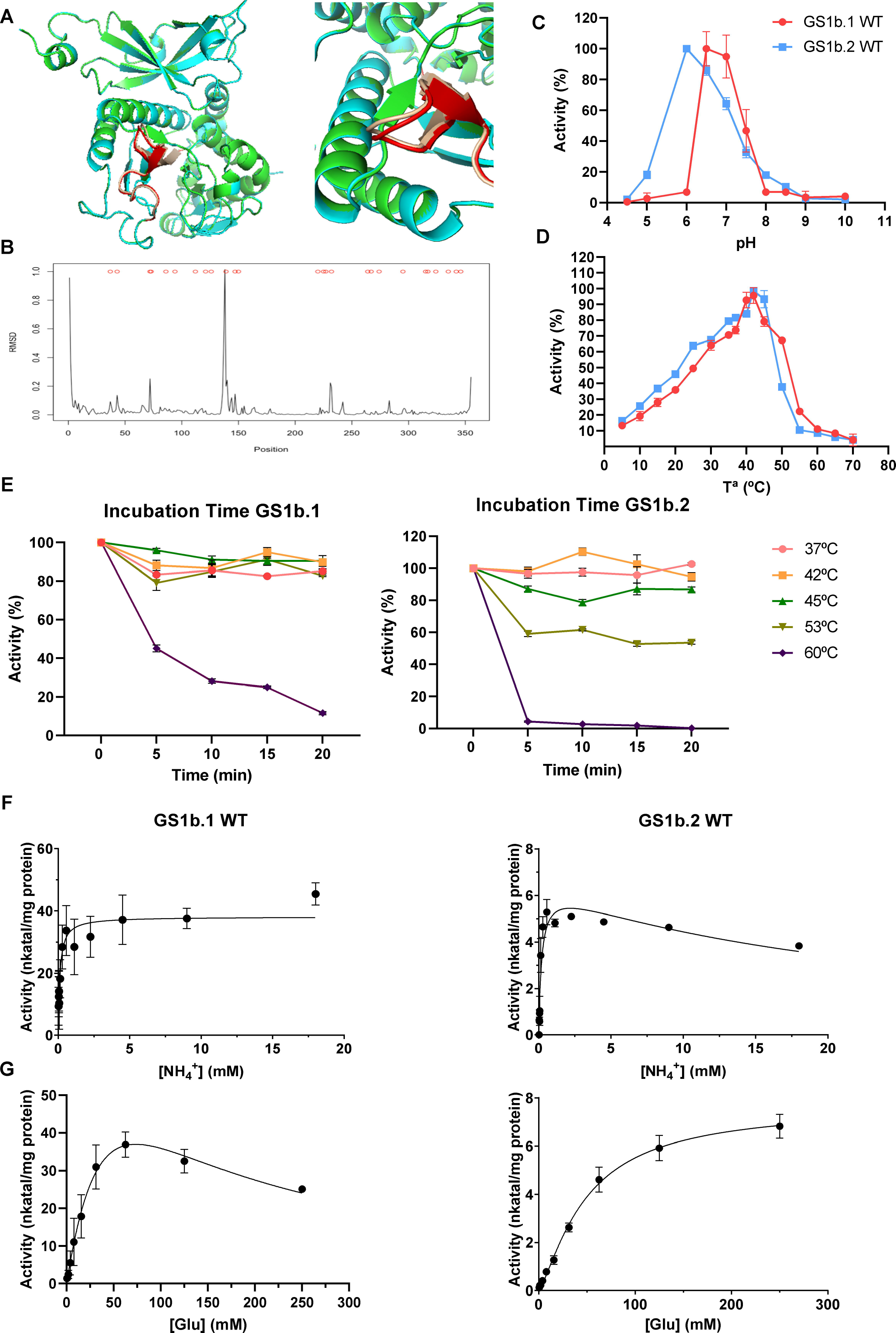
Enzymatic characterization of recombinant GS1b.1 and GS1b.2 isoforms. **A.** Comparison of GS1b.1 and GS1b.2 subunit structure GSb1.1 is represented in green and GS1b.2 in cyan. The region that presented most differences between GS1b.1 and GS1b.2 (amino acids from 125 to 150) are represented in red and pink respectively. **B.** Root mean square deviation (RMSD) values between GS1b.1 and GS1b.2 monomers structure. **C.** Enzyme activity at different assay pH (from 4.5 to 10) for GS1b.1 (red line) and GS1b.2 (blue line). **D.** Enzyme activity at different assay temperature (from 5 to 70°C) for GS1b.1 (red line) and GS1b.2 (blue line). **E.** Thermal stability of GS1b.1 and GSb1.2 at different temperatures (37, 42, 45, 53 and 60°C) after different preincubation times (from 0 to 20 min). **F.** Kinetics of GS1b.1 and GS1b.2 for ammonium. **G.** Kinetics of GS1b.1 and GS1b.2 for glutamate. Error bars show the SD. Mean values are composed with at least three independent determinations.

**Table 1.**
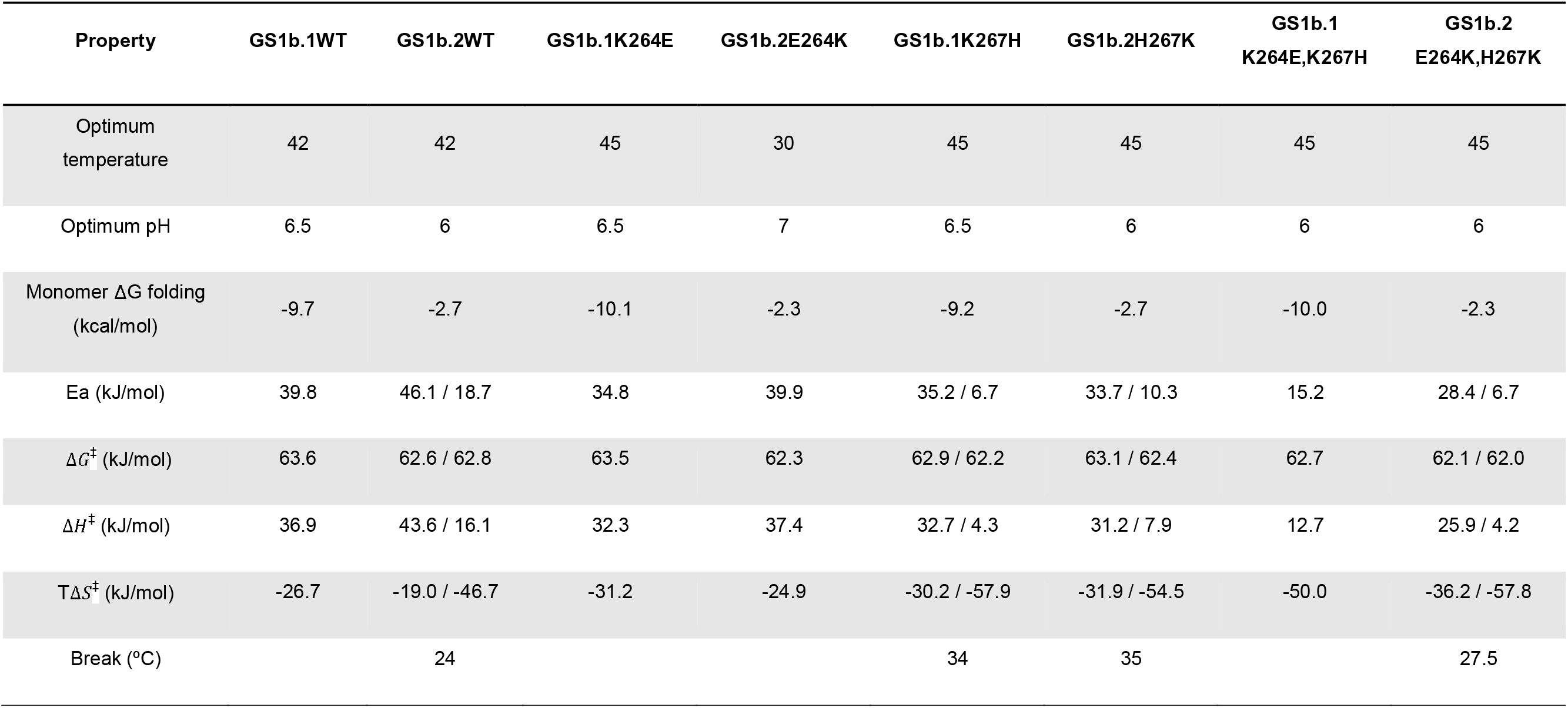
Physicochemical properties of the wild type PpGS1b.1 and PpGS1b.2 and their mutated versions.

**Table 2.** Kinetic properties of the wild type PpGS1b.1 and PpGS1b.2 and their mutated versions.

| Substrate | GS1b.1WT | GS1b.2WT | GS1b.1K264E | GS1b.2E264K | GS1b.1K267H | GS1b.2H267K | GS1b.1<br>K264E,K267H | GS1b.2<br>E264K,H267K |
| --- | --- | --- | --- | --- | --- | --- | --- | --- |
| NH<sub>4</sub><sup>+</sup> | Vmax 38.15 | Vmax 6.48 | Vmax 21.01 | Vmax 16.36 | Vmax 79.3 | Vmax 58.26 | Vmax 36.05 | Vmax 56.88 |
|  | Km 0.12 | Km 0.21 | Km 0.16 | Km 0.16 | Km 0.06 | Km 0.05 | Km 0.02 | Km 0.09 |
|  |  | Ki 22.57 |  |  | Ki 13.14 |  |  |  |
| Glu | Vmax 101.6 | Vmax 7.66 | Vmax 16.82 | Vmax 15.69 | Non saturated | Non saturated | Non saturated | Non saturated |
|  | Km 64.18 | nH 1.311 | Km 2.20 | Km 18.41 |  |  |  |  |
|  | Ki 84.51 | EC50 48.63 |  |  |  |  |  |  |
| Mg<sup>2+</sup> | Vmax 71.32 | Vmax 5.64 | Non saturated | Vmax 25.43 | Non saturated | Non saturated | Non saturated | Non saturated |
|  | nH 1.66 | nH 2.33 |  | nH 1.61 |  |  |  |  |
|  | EC50 14.49 | EC50 10.87 |  | EC50 26.78 |  |  |  |  |
| ATP | Vmax 24.96 | Vmax 7.39 | Vmax 29.73 | Vmax 27.40 | Vmax 59.82 | Vmax 28.57 | Vmax 38.69 | Vmax 100.6 |
|  | Km 0.18 | Km 0.29 | Km 0.11 | Km 0.21 | Km 0.06 | Km 0.39 | Km 0.09 | Km 0.42 |
|  | Ki 5.88 |  | Ki 3.19 | Ki 8.23 | Ki 5.85 | Ki 8.76 |  | Ki 5.06 |

EC50 in mM; Ki in mM; Km in mM; nH is dimensionless; Vmax in nkat/mg protein. EC50, the concentration of substrate that produces a half-maximal reaction rate; Ki, dissociation constant for substrate binding; Km, Michaelis-Menten constant; nH, Hill slope; Vmax, maximal reaction rate.

Both isoforms were tested over a wide pH range; GS1b.1 maximum activity was reached at pH 6.5 while that of GS1b.2 maximum activity was reached at pH 6 (Figure 5C). The activity of both enzymes increased with the reaction temperature, reaching the maximum activity at 42 °C (Figure 5D). These data have allowed the calculation of the activation energy (Ea) for each enzyme (Table 1). The Ea was different for both enzymes: the Ea of GS1b.1 was 39.9 kJ/mol, and the Ea values of GS1b.2 for its elemental reaction steps were 46.1 kJ/mol and 18.7 kJ/mol, with a break point at 24 °C. Regarding the thermal stability, GS1b.1 was very stable, only decreasing its activity at 60 °C after 5 minutes of preincubation, although it never completely lost its activity, even after 20 min at 60 °C (Figure 5E). However, GS1b.2 showed a decreased activity even after 5 minutes of preincubation at 45 °C with almost a total loss of activity after 5 minutes at 60 °C (Figure 5E).

GS1b.1 and GS1b.2 showed distinctive behaviors for ammonium and glutamate (Figure 5F,G). GS1b.2 exhibited substrate inhibition for ammonium (K_i_ 22.57 mM). The affinities of both enzymes for ammonium were high (GS1b.1 K_m_ 0.12 mM and GS1b.2 K_m_ 0.21 mM). However, the V_max_ was 5.88 times higher for GS1b.1 (Table 2). Regarding to glutamate, GS1b.1 showed substrate inhibition at high concentrations (K_i_ 84.51 mM), while GS1b.2 presented positive cooperativity. In both cases, the affinity was very low (GS1b.1 K_m_ 64.15 mM and GS1b.2 EC50 48.63 mM), with large differences in the V_max_ values of both enzymes (GS1b.1 101.6 nkat/mg protein and GS1b.2 7.66 nkat/mg protein) (Table 2). GS1b.1 and GS1b.2 showed equal behavior for Mg_2_^+^, with positive cooperativity and similar affinity (EC50 values of 14.49 and 10.87 mM, respectively) but different V_max_ values (71.32 and 5.64 nkat/mg protein, respectively) (Figure S4, Table 2). Finally, the affinities for ATP were high and similar for both enzymes (K_m_ of 0.18 and 0.29 mM for GS1b.1 and GS1b.2, respectively), with a higher V_max_ for GS1b.1 (24.96 nkat/mg protein) than for GS1b.2 (7.39 nkat/mg protein). However, there was substrate inhibition for GS1b.1 at moderate levels of ATP (K_i_ 5.88 mM).

### Analysis of mutant proteins

To determine the roles that certain residues could play in GS activity, mutants of GS1b.1 and GS1b.2 were obtained by exchanging amino acids at positions 264 and 267. These residues belong to a region that accumulates a significant number of differences between the two isoforms and is important for stability, as shown by the *in silico* substitution analysis (Figure S3). Additionally, these residues have been selected based on their charge and structural differences between both GSs. The amino acid swapping at positions 264 and 267 seemed to produce only slight changes in the subunit arrangement, even in the double mutant. Calculation of hydrogen bonds revealed interactions between residues 264 and 267 with those present at positions 261, 263, 265 and 268. These residues were analyzed in detail, and only small differences in their arrangements could be observed (Figure 6A-G, S5). The quaternary structures of the mutants also showed no significant differences when compared (Figure S6) and the thermodynamic stability of the monomers was similar to that of the WT (Table 1).

**Figure 6.**
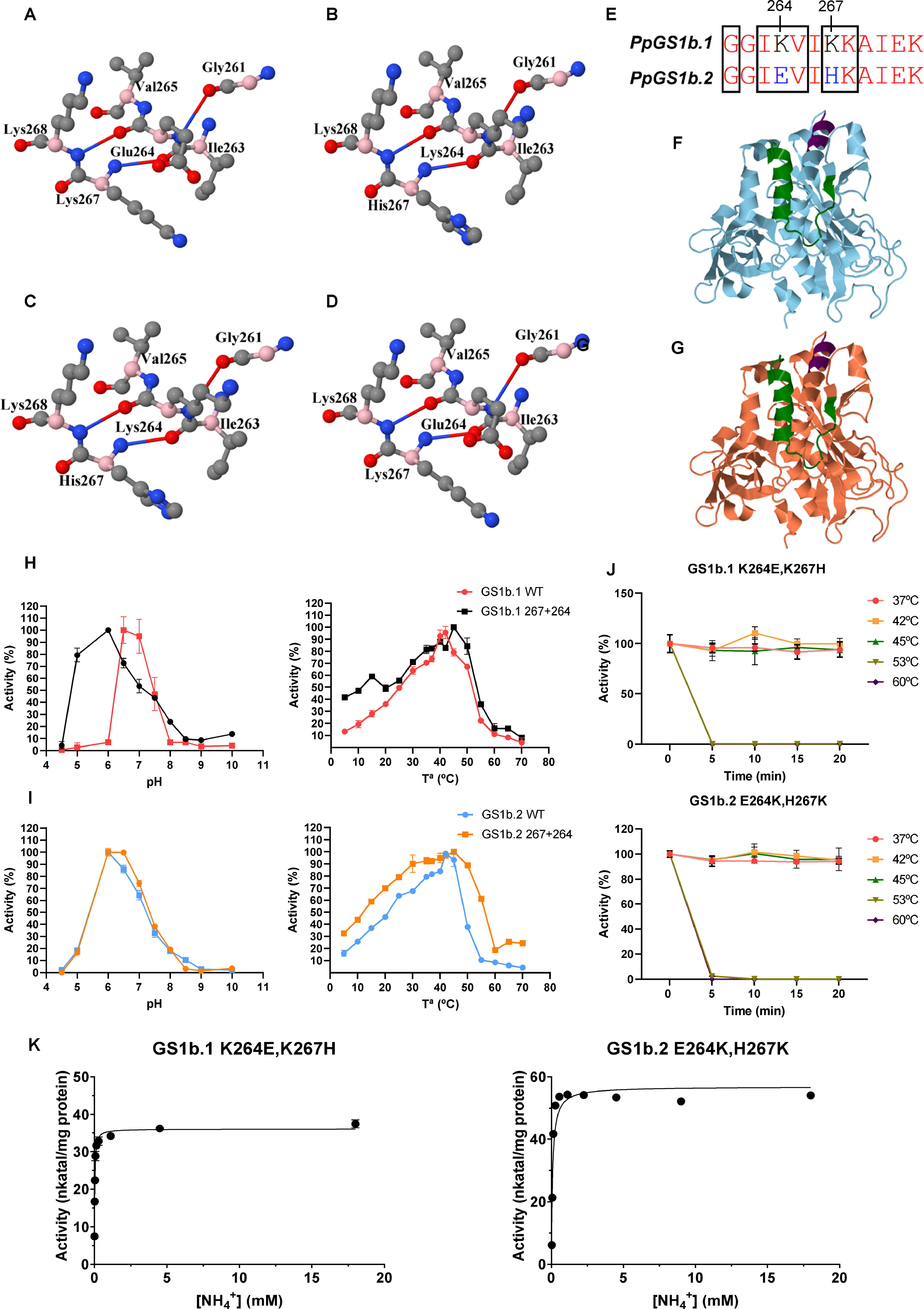
Characterization of mutated GS1b.1 and GS1b.2 proteins. Disposition of the amino acids, either those that have been exchanged and those associated with them by hydrogen bonds in the GS1b.1 K264E (**A**), GS1b.2 E264K (**B**), GS1b.1 K267H (**C**) and GS1b.2 H267K (**D**) mutants. Alpha carbons of the amino acids are represented in pink. **E**. Amino acid region affected by mutations. Subunit structure of the GS1b.1 (**F**) and GS1b.2 (**G**) double mutant. Amino acids exchanged and amino acids associated with them by hydrogen bonds are represented in dark magenta. Amino acids from 330 to the end of the protein are represented in green. **H**. Comparison of the physicochemical properties of the GS1b.1 WT and its double mutant. **I**. Comparison of the physicochemical properties of the GS1b.2 WT and its double mutant. **J**. Thermal stability of the double mutants at different temperatures (37, 42, 45, 53 and 60°C) after different preincubation times (from 0 to 20 min). **K.** Kinetics of GS1b.1 and GS1b.2 double mutants for ammonium. Error bars show the SD. Mean values are composed with at least three independent determinations.

Compared to WT, none of the optimal pH values were affected in any of the mutants tested, except for GS1b.2E264K, where the optimum was reached at pH 7 (Figure S7A), and the double mutants, where the optimum pH was 6 for both enzymes (Figure 6H, S7).

A slight increase in the optimal temperature (45 °C) was detected in all mutants except for GS1b.2E264K, which experienced a large change in its optimal temperature (30 °C) (Figure 6I, S7B). Although the activity patterns in response to reaction temperature were similar in the mutants with respect to the WT enzymes, the activity was slightly higher at all temperatures in the GS1b.1K267H single and GS1b.2 double mutants. In the case of GS1b.1 K264E and GS1b.2 H267K, the activity was higher at temperatures above the optimum (45 °C). Finally, the GS1b.1 double mutant retains considerable activity levels (>40%) even at very low reaction temperatures, such as 4 °C (Figure 6I, S7B). Ea was barely affected (Table 1) in GS1b.1K264E (34.8 kJ/mol). In contrast, the GS1b.1 double mutant Ea was strongly affected (15.2 kJ/mol), and GS1b.1 K267H showed different Ea values for its elemental reaction steps (35.2 kJ/mol and 6.7 kJ/mol), similar to GS1b.2 WT. However, GS1b.2 E264K presented a unique Ea for its reaction (39.9 kJ/mol), and different Ea values were detected for the elemental reaction steps of GS1b.2 H267K (33.7 kJ/mol and 10.3 kJ/mol) and the GS1b.2 double mutant (28.4 kJ/mol and 6.7 kJ/mol). Interestingly, all GS1b.1 mutants experienced decreases in their thermostability compared to that of the WT, and only GS1b.2H267K showed an increased thermostability compared to GS1b.2 WT (Figure 6J, S7C).

GS1b.1 behavior regarding ammonium was only modified in the GS1b.1K267H mutant, which showed substrate inhibition for ammonium (K_i_ 13.14 mM). Furthermore, the affinity was increased in this mutant, GS1b.2H267K, and both double mutants (K_m_ between 0.02 and 0.09 mM). Meanwhile, all the GS1b.2 mutants lost substrate inhibition by ammonium, and all exhibited normal hyperbolic saturation (Figure 6K, S8, Table 2). Regarding glutamate, GS1b.1K264E lost substrate inhibition, now presenting normal hyperbolic saturation with an increase in its affinity (K_m_ 2.2 mM) accompanied by a reduction in V_max_ (16.82 nkat/mg protein). Additionally, none of the mutants in 267 and double mutants reached saturation and seemed to have lost affinity for this substrate, as occurred with Mg_2_^+^ in all the mutants except for GS1b.2E264K (Figure S9, S10, Table 2). GS1b.1 mutants exhibited substrate inhibition by ATP, but only the double mutants of GS1b.1 lost substrate inhibition by ATP and presented a normal hyperbolic saturation for this substrate (Figure S11, Table 2). Interestingly, all GS1b.2 mutants presented inhibition by ATP (K_i_ ranging from 5.06 to 8.76 mM), in contrast to the hyperbolic Michaelis-Menten saturation exhibited by the WT (Figure S11, Table 2).

## DISCUSSION

The phylogenetic analysis carried out in this work (Figure 1) grouped the new GS isoform (GS1b.2) within the conifer GS1b.1 group. Furthermore, the identification of *GS1b.2* in the genome, its different promoter sequences, including different TF binding sites (Figure S1B), and its different gene expression patterns rule out the possibility that it is an allelic variant of *PpGS1b.1* (HF548531.1), suggesting that *PpGS1b.2* (KU641799.1; KU641800.1) is likely the result of a gene duplication. The presence of *GS1b.2* in members of the genera *Pinus* and *Picea* indicates (Figure 1) that this gene duplication should have taken place in a common ancestor of these two groups but not of the entire *Pinaceae* family since orthologs of *GS1b.2* have not been identified in other conifers. Gene duplication is very common in plants (De Smet and Van de Peer, 2012), and it could lead to the acquisition of new functions (neofunctionalization) or simply to redundant activity to maintain the correct metabolic flux, as occurs with GS in *Populus* and rice (Yamaya and Kusano, 2014; Castro-Rodríguez *et al*., 2015), contributing to metabolic homeostasis (Moreira *et al*., 2022). In fact, the GS1b family in angiosperms has been extended by gene duplication so that different isoenzymes can play nonredundant or synergistic roles within the plant, as proposed for *Arabidopsis GS1* genes (Ji *et al*., 2019).

To explore the possible neofunctionalization of this new *GS* after gene duplication, the expression patterns of the maritime pine *GS* genes were analyzed in different organs and tissues (Figure 2–4). *PpGS1b.2* appears to be expressed primarily in developing organs and tissues and is tightly regulated throughout embryonic development. This contrasts with *PpGS1b.1* expression, which was high in all analyzed samples. This could indicate a strong regulation of *PpGS1b.2* at both the localization and expression levels, suggesting a specialized function. The expression of *PpGS1b.2* is consistent with the association of some *GS1b* isogenes with plant developmental processes in angiosperms (Habash *et al*., 2001; Tabuchi *et al*., 2005; Martin *et al*., 2006; Lothier *et al*., 2011; Funayama *et al*., 2013; Goodall *et al*., 2013; Bao *et al*., 2014; Guan *et al*., 2015; Urriola and Rathore, 2015; Gao *et al*., 2019; Ji *et al*., 2019; Wei *et al*., 2021; Fujita *et al*., 2022). These data suggest an evolutionary convergence that has led to the emergence of GS1b isoforms with similar roles in different plant species. The expansion of the GS1b family in certain conifers supports that GS1b diversification in angiosperms responds to different plant needs associated with N assimilation (Hirel and Krapp, 2021). In pine, *GS1b.1* has also been associated with this function due to its expression during zygotic and somatic embryo development (Pérez-Rodríguez *et al*., 2005). All these expression data pose different hypotheses about the role of this new isoenzyme: a) GS1b.2 could support GS1b.1 activity in developing tissues with a high demand for glutamine or assimilated N; and b) GS1b.2 could play a specific role in certain developing tissues.

To explore the differential roles of GS1b.1 and GS1b.2 in maritime pine, the structure, as well as the physicochemical and kinetic properties of both enzymes, were analyzed. Modeling of both maritime pine GS1b isoforms reports small differences between GS1b.1 and GS1b.2 when their tertiary and quaternary structures were compared (Figure 5A,B; S2). However, any minor difference in subunit arrangements could be of great importance since the GS active site is formed by the N- and C-terminal domains of adjacent subunits (Llorca *et al*., 2006).

Although GS1b.1 and GS1b.2 are very similar in their primary sequences and structures, quite a few differences have been found in their properties. The thermodynamic stability of GS1b.1 was three times higher than that shown by GS1b.2 (Table 1). Both isoenzymes present similar values (approximately 63 kJ/mol) for the change in Gibbs free activation energy (Δ*G*^‡^), but their kinetic response to temperature changes below and above 24°C may be very different (Table 1). For the new isoform, Δ*G*^‡^ and the rate-limiting step are dominated by different activation parameters at different operating temperatures: Δ*H*^‡^ for temperatures below 24°C and TΔ*S*^‡^ for temperatures above 24°C. In contrast, GS1b.1 showed a nonvariable activation energy throughout the whole range of temperatures assayed (Table 1). These differences in dominant activation parameters could reflect functional differences between the two active sites, as has been previously suggested for glutamine synthetase isoforms from other sources (Wedler and Horn, 1976).

The optimum pH levels for GS1b.1 and GS1.b2 are 6.5 and 6, respectively (Table 1; Figure 5C), similar to those of GS1b.2 and GS1b.3 from poplar (Castro-Rodríguez *et al*., 2015). Interestingly, these optimal pH values are lower than the cytosolic pH (7.1-7.5) (Zhou *et al*., 2021), which could be a mechanism to avoid enzyme inhibition by the acidification process associated with GS activity and ammonium (Hachiya *et al*., 2021). The optimum temperature for both enzymes (42 °C) (Table 1; Figure 5D) is very similar to that shown by GS1b isoenzymes in other plants (Zhao *et al*., 2014, Castro-Rodríguez *et al*., 2015). However, both GS1b enzymes had exceptional thermostability compared to other GS1b enzymes of plants (Figure 5E) (Sakakibara *et al*., 1996; Zhao *et al*., 2014; Castro-Rodríguez *et al*., 2015). Concerning glutamate and ATP, GS1b.1 exhibited substrate inhibition behavior, as previously observed for *Arabidopsis* GLN1;3 (Table 2; Figure 5G) (Ishiyama *et al*., 2004b). These inhibitions are consistent with the role of GS1b.1 in primary nitrogen assimilation in pine and its high expression since high levels of glutamate and ATP, outside of their homeostatic ranges, could indicate metabolic and energetic problems in the cell that may result in unnecessary or detrimental large-scale nitrogen assimilation. Interestingly, GS1b.2 exhibited positive cooperativity for glutamate (Table 2; Figure 5G) and showed substrate inhibition for ammonium (Table 2; Figure 5F). The positive cooperativity mechanism provides high sensitivity to fluctuating substrate concentrations (Levitzki and Koshland, 1976), enabling GS1b.2 to respond rapidly to changes in glutamate availability. In this case, the inhibition of GS1b.2 by ammonium could lead to control of the levels of the final product or to a specific function on the signaling pathway of one of its substrates. This is because both the end product and the substrate of the GS/GOGAT cycle, glutamate and ammonium, have been reported to play roles in plant growth and development (Qiu *et al*., 2020; Ortigosa *et al*., 2021), where GS could act as an integrating link for both signaling pathways. Interestingly, glutamate has been described to play important roles in seed germination (Kong *et al.*, 2015), root architecture (Forde, 2014; López-Bucio *et al.*, 2019) and pollen germination and pollen tube growth (Michard *et al.*, 2011; Wudick *et al.*, 2018), among other functions (Qiu *et al.*, 2020). Ammonium has been shown recently to modulate plant root architecture in pine seedlings (Ortigosa *et al.*, 2022). Therefore, based on the *PpGS1b.2* expression patterns, the kinetic characteristics toward glutamate, and previous works, this enzyme could be involved in developmental processes. Furthermore, this could also be a mechanism to avoid high GS activity levels when ammonium is in excess, which could lead to excessive cytosol acidification (Hachiya *et al.*, 2021) of sensitive cells in developing tissues.

The structural, physicochemical, and kinetic analysis carried out in this work on the mutant enzymes showed some differences from the WT isoforms, but almost none of them achieved a complete exchange of the properties between GS1b.1 and GS1b.2. The mutations tested in this work did not greatly affect the protein structure, either in the surroundings of the exchanged amino acids and the subunit structure (Figure 6A-G) or in the quaternary structure (Figure S6), which could explain why the thermodynamic stability of the mutants was not compromised in any case (Table 1). Although all the mutants presented alterations in the activity levels at the different pH values and temperatures analyzed in comparison with the WT, only GS1b.1K264E,K267H and GS1b.2E264K produced variations in the optimal pH, and only GS1b.2E264K presented a considerable variation in its optimum temperature (Figure S7B) and Ea (Table 1). In fact, among all the mutants, GS1b.2E264K presented the greatest number of changes in physicochemical properties. In fact, this could indicate that none of these amino acids have strong involvement in these enzyme properties or, perhaps, that the changes that can produce these mutations are being buffered by other residues.

Interestingly, these mutations had large effects on the kinetic properties (Table 2; Figure 6K; S8-S11). The results suggest that these residues are involved in ammonium affinity. Although it has been described that the presence of glutamine and serine at positions 49 and 174, respectively, is essential for the high affinity for ammonium in *Arabidopsis* GS (Ishiyama *et al*., 2006), these residues are not present in either GS1b.1 or GSb1.2 of *P. pinaster*. Previous kinetic studies have shown the presence of high-affinity GS isoforms that either do not have this combination of amino acids or have none of them (Sakakibara *et al*., 1996; de la Torre *et al*., 2002; Yadav, 2009; Zhao *et al*., 2014; Castro-Rodríguez *et al*., 2015). These previous works and the current results support the hypothesis proposed by Castro-Rodríguez *et al*. (2015), indicating that key residues determining GS behavior for ammonium may vary between plant species.

Mutations have produced a great number of changes in the behavior of these enzymes against their substrates and in their kinetic parameters. However, a reversal has only been achieved for ATP in double mutants, suggesting that the differences in these properties are due to the collaborative efforts of several residues, probably those that differ between the two enzymes. This may indicate that GS1b.1 and GS1b.2 have undergone evolutionary selection so that the two enzymes satisfy different plant needs, with only minor changes in their amino acid sequences. This hypothesis is also supported by the differences between the two enzymes at the structural stability level (Figure S3). When introduced at certain positions, some amino acids had a large effect on the protein stability of one isoform but not the other. The region between amino acids 260-300 of GS1b.2 was particularly affected by the introduction of some amino acids, but none of these substitutions appear to produce similar effects on GS1b.1. In fact, these data suggest that the two enzymes are probably undergoing different evolutionary paths.

## EXPERIMENTAL PROCEDURES

### Sequence identification and phylogenetic analyses

The phylogenetic analysis was made using protein sequences of plant GS that were obtained from online public databases or assembled from transcriptomic data contained in the SRA database at the NCBI except for *Pinus pinaster* sequences that were cloned and sequenced in the present work (Table ◻ S1). For the sequence obtaining, the procedure presented in Valderrama-Martin *et al*. (2022) was followed. Briefly, *tblastn* was used in BLAST searches (Altschul *et* ◻ *al*., ◻ 1990) using GS1b.1 from *Pinus taeda* as the query. Transcriptomic assemblies were made in the web platform Galaxy (Afgan *et* ◻ *al*., ◻ 2018). Raw reads were trimmed using *trimmomatic* (Bolger *et* ◻ *al*., ◻ 2014) and assembled with *Trinity* (Grabherr *et* ◻ *al*., ◻ 2011). Database identifiers, names and species for the different GS sequences are presented in Table ◻ S1. All protein sequences used in the present work are available in Dataset S1.

The sequence data set was composed of 96 GS proteins. The phylogenetic analysis was mainly focused on conifer GS sequences. The alignment and phylogenetic analysis were conducted as described in Valderrama-Martin *et al*. (2022) using MEGA version ◻ 11 (Tamura *et* ◻ *al*., ◻ 2021). The alignment was conducted with *muscle* (Edgar, ◻ 2004). The phylogenetic analysis was carried out through a maximum-likelihood estimation with complete deletion of gaps, the missing data, and the Jones–Taylor–Thornton amino acid substitution model (Jones *et* ◻ *al*., ◻ 1992). Nearest-neighbor interchange was used for tree inference. The initial tree was constructed using the NJ/BioNJ method. The phylogeny test was performed using the bootstrap method with 1000 replications. The GS sequences of *Chlamydomonas reinhardtii* were used as outer group. The distance matrix and the original tree in Newick format are available in Datasets S2 and S3. The original tree was visualized with the Interactive Tree of Life web tool (Letunic and Bork, ◻ 2019).

### Protein structure prediction and modeling

For the 3D modeling and structure predictions of *P. pinaster* GS1b.1 and GS1b.2 individual subunits, Alphafold (Jumper *et al*., 2021; Varadi *et al*., 2022) through ColabFold (Mirdita *et al*., 2022) has been used. ColabFold allows faster protein structure prediction by integrating MMseqs2 for multiple sequence alignments and AlphaFold2, but it does not allow the structure prediction of large protein subunits or complexes. The quaternary structure prediction has been achieved using Alphafold’s models as input for the Galaxy Package, a combination of several programs that have been designed based on sequence and structure information together with physical chemistry principles (Shin *et al*., 2014). The models obtained from ColabFold were employed for the comparison and graphic representation of the protein structure in PyMOL (https://pymol.org/2/) and in Jmol (http://www.jmol.org/). Jmol was also used for the calculation of the hydrogen bonds. Quaternary structure models obtained with Alphafold and the Galaxy Package has been used in PyMol for the structure analysis and comparison of the models. The thermodynamic stability of the monomers has been determined using models obtained in AlphaFold together with the “foldx.mut()” function of the “ptm” R package (Aledo, 2021).

### Plant material

Maritime pine seeds (*P. pinaster* Aiton) from *Sierra Segura y Alcaraz* (Albacete, Spain) (ES17, Ident. 09/10) were provided by the *Red de Centros Nacionales de Recursos Genéticos Forestales* of the Spanish *Ministerio para la Transición Ecológica y el Reto Demográfico* with the authorization number ESNC103. Pine seeds were imbibed for 48 h in water with aeration to induce germination. Seeds were germinated in vermiculite. Seedlings were grown in plant growth chambers (Aralab Fitoclima 1200, Rio de Mouro, Portugal) under 16 ◻h light photoperiod, a light intensity of 125 ◻ μmol ◻ m^−2^ ◻ s^−1^, a constant temparature of 23 ◻ °C, 50% relative humidity and watered twice a week with distilled water. Embryo and seedling samples were harvested at different stages: dry, post-imbibition and germinated (0.5 cm of emerged radicle) embryos; and one-week-old from emergence (Stage 1) and one-month-old from emergence seedlings (Stage 2). At the harvest, seedlings were divided into their different organs. For the measure of *GS* gene expression in different sections of roots, 2 months-old seedlings were used. The samples were immediately frozen in liquid N and stored at −80 °C until powdering with a mixer mill MM400 (Retsh, Haan, Germany) and further analyses were conducted.

Plant material and cDNA to analyze *GS* gene expression levels in maritime pine tissues from one-month-old seedlings were previously obtained by Cañas *et al*. (2017). RNA samples from 14 tissues isolated through laser capture microdissection were employed. The cDNA was synthesized and amplified as described by Cañas *et al*. (2014).

Samples from Cañas *et al*. (2015) were used to analyze *GS* gene expression in needles of adult trees. Briefly, needle whorls corresponding to the annual growth of a single year were harvested from different 25 years old *P. pinaster* specimens at *Los Reales de Sierra Bermeja* (Estepona, Spain). Whorls were named from 0 to 3 referring to the year of appearance of that whorl. Whorl 0 was first collected in May when it was completely formed. Samples were collected each month throughout 2012, were immediately frozen in liquid nitrogen and stored at −80°C until their utilization for RNA extraction. Buds and nascent needles were collected from the same adult specimens once a week during April of 2013. For gene expression analyses three different trees were employed.

Juvenile and mature phloem, together with male and female strobili were harvested from 25 to 35-year-old maritime pines located *at Los Reales de Sierra Bermeja* (Estepona, Spain) Juvenile xylems were collected from the last 5 internodes in the crown and mature xylem from the base of the trunk of 28 to 31-year-old maritime pines from *Los Reales de Sierra Bermeja* by removing bark and phloem and scraping with a sterile blade. (Villalobos, 2008). All the tissues were frozen immediately using liquid nitrogen and storage at −80 °C until use

Zygotic embryos from *P. pinaster* were obtained from a single maritime pine seed orchard (PP-VG-014, Picard, Saint-Laurent-Médoc, France) and collected at different developmental stages (Avila et al., 2022). All samples were frozen in liquid nitrogen and stored at −80°C until use.

### RNA extraction and RT-qPCR

Total RNA from maritime pine samples was extracted following Canales et al. (2012). RNA concentration and purity (A260/A280) was then quantified using a NanoDrop© ND-1000 spectrophotometer (ThermoFisher Scientific, Walthman, MA, USA). The integrity of the RNA was checked by electrophoresis. iScript Reverse Transcription Supermix (Bio-Rad, Hercules, CA, USA) was used for the reverse transcription of 500 ng of total RNA of each sample in a final volume reaction of 10μL including 2μL of reaction buffer and 0,5μL of reverse transcriptase enzyme with the following conditions in a thermal cycler with the following conditions: 30 min at 42°C; 10 min at 65°C; hold at 4°C.

For the RT-qPCR analysis, three biological samples were used with three technical replicates each. The qPCR was carried out using 5 μL SsoFastTM EvaGreen® Supermix (Bio-Rad, Hercules, CA, USA), 10 ng of cDNA, 20 pmol of each primer in a total reaction volume of 10 μL on a C1000^™^ Thermal Cycler with a CFX384^™^ Touch Realm-Time PCR Detection System (Bio-Rad, Hercules, CA, USA) with the following conditions: initial denaturation step at 95°C 2 min; 40 cycles of denaturation at 95 °C 5 s and elongation at 60 °C 20 s. Finally, a melt curve was developed from 65 to 95 °C with increments of 0.5 °C each 5 s. Two maritime pine *saposin-like aspartyl protease* and *RNA binding protein* genes were used as reference for results normalization (Granados et al., 2016). Expression data have been analyzed using the *qpcR* R library and the MAK3 model (Ritz and Spiess, ◻ 2008). The primers used for RT-qPCR assays are presented in Table S2.

### Cloning, mutagenesis, recombinant expression, and purification of GS1b.1 and GS1b.2

In the search for new *GS* genes in conifers, 3 genes in *P. pinaster* have been identified in transcriptome databases that were named as *PpGS1a*, *PpGS1b.1* and *PpGS1b.2*. The cDNA of the three genes were amplified by PCR using iProof HF Master Mix (Bio-Rad, Hercules, CA, USA) and cloned into the pJET1.2 vector (ThermoFisher Scientific, Walthman, MA, USA) following the manufacturers’ instructions. The used primers were designed from sequences obtained from the maritime pine transcriptome assembled in Cañas *et al*. (2017). Primers are shown in Table S2. *PpGS1a* was obtained from amplified cDNA of emerging needles (EN) isolated in Cañas *et al*. (2017). *PpGS1b.1* and *PpGS1b.2* were obtained from amplified cDNA of developing root cortex (DRC) isolated in Cañas *et al*. (2017).

For protein recombinant expression, the CDS of wild type (WT) *PpGS1b.1* and *PpGS1b.2* were subcloned into pET30a vector (Merck, Darmstadt, Germany) including a N-terminal 6xHis-tag by PCR. For this task, AseI and XhoI sites were added to *PpGS1b.1* 5′and 3′ends respectively while NdeI and XhoI sites were added to *PpGS1b.2* 5′and 3′endings respectively. These restriction sites along with the 6xHis-tag were introduced by PCR. Used primers are listed in Table S2. The plasmid and PCR product were then cut using the appropriate restriction enzymes and the PCR product was inserted into the plasmid using T4 DNA ligase.

Plasmids were transformed and expressed in the *Escherichia coli* strain BL-21 (DE3) RIL cells (Agilent, Santa Clara, CA, USA). For protein expressions, the bacterial clones were grown at 37 °C and 180 rpm in an orbital shaker with 500 mL of Luria-Bertani medium supplemented with kanamycin (0.05 mg/mL) and chloramphenicol (0.034 mg/mL). When the optical density (OD) reached a 0.5-0.6 value at 600 nm, cultures were tempered and isopropyl-β-D-thiogalactoside was added to a final concentration of 1 mM to induce protein expression. Once the isopropyl-β-D-thiogalactoside was supplied the cultures were incubated at 25 °C and 120 rpm for 5 hours, the cells were collected by centrifugation. The bacterial pellet was resuspended in 5 mL of buffer A (Tris 50 mM pH 8; NaCl 300 mM; imidazole 250 mM) with 4 mg of lysozyme and incubated for 30 min in ice, bacteria were then lysed by ultrasonication with 20 pulses of 5 seconds at 20% amplitude with 5 seconds rest between pulses in a Branson Sonifier® Digital SFX 550 (Branson Ultrasonics, CT, USA). The soluble fraction was clarified by centrifugation (1620 x g at 4 °C for 30 min). Proteins from the soluble fraction were purified by affinity chromatography with Protino Ni-TED PackedColumns2000 (Macherey-Nagel, Düren, Germany) based on the His-tag tail. The soluble fraction from bacterial lysate was loaded in a column previously equilibrated with buffer A. Protein elution was performed by adding buffer B (Tris 50 mM pH8; NaCl 300 mM; imidazole 250 mM) and a total of 9 mL of eluate was recovered in 1 mL fractions. Collected fractions were quantified by Bradford (Bradford, 1976) and analyzed on SDS-page and western-blot using GS-specific antibodies obtained from rabbit (Figure S12) (Cantón *et al*., 1996). Fractions containing the proteins were concentrated with Amicon® Ultra-15 Centrifugal Filters Ultracel®-100K (Merck-Millipore, Burlington, Massachusetts, State of Virginia) with 100 kDa pores and the resulting concentrate was stored in 50% (v/v) glycerol at −20 °C for later kinetic measurements and physicochemical analyses.

### Site-directed mutagenesis

Considering characteristics and properties of differing amino acids between GS1b.1 and GS1b.2, residues at position 264 and 267 were selected to be shifted between both isoenzymes. Site-directed mutagenesis was carried out following Edelheit *et al*. (2009). The wild type CDS from those sequences included on the pET30a vector were amplified by PCR using two reverse-complementary primers (Table S2) that already included the mutation to be introduced. The primers were used separately in a PCR reaction using 50 or 500ng of plasmid and 10 pmol of each primer. The final products of both reactions were then mixed and hybridized. The PCR products were checked out in an agarose gel and purified using NucleoSpin® Gel and PCR Clean-up (Macherey-Nagel, Düren, Germany). Finally, the PCR product was digested with FastDigest® DpnI (ThermoFisher Scientific, Walthman, MA, USA) to degrade the vector used as template for the amplification.

### Physicochemical assays

Physicochemical properties were determined by conducting the transferase assay as described in Cánovas *et al*. (1991). Reactions were carried out in 96 well microtiter plates with a final reaction volume of 150 μL. The reaction mix contained 90.6 mM MOPS pH 7, 20 mM arsenate, 2.93 mM MnCl_2_, 60 mM NH_2_OH and 0.4 mM ADP. When determining the optimal pH level for the activity of the different isoforms, different buffers were used instead when determining the optimal pH level for the activity of the different isoforms: acetate (4.5-5); MES (6-6.5); HEPES (7-7.5); Tris (8-8.5); and sodium carbonate (9-10). The reaction was initiated by adding glutamine in a final concentration of 120 mM and, after 15 minutes of incubation at 37 °C, 150 μ of STOP solution (10% FeCl_3_ · 6 H_2_O in HCl 0.2 N; 24% trichloroacetic acid and 5% HCl) was added to stop the reaction. Finally, the plate was centrifugated for 3 minutes at 3220 x g and 100 μL of the reaction volume were withdrawn for its absorbance measurement at 540 nm in a PowerWave HY (BioTek, Winooski, VT; USA) plate lector. For thermostability characterization, proteins were preincubated at different times and temperatures before adding the reaction mix.

### Kinetic assays

For the quantification of the kinetic properties, biosynthetic assays were carried out as described by Gawronski and Benson (2004) with some modifications. Reactions were conducted in 96 wells microtiter plates in a final volume of 100 μL. GS activity was determined as a function of NADH absorbance depletion at 340 nm in a coupled reaction using lactate-dehydrogenase (LDH, EC 1.1.1.27) and pyruvate kinase (PyrK, EC 2.7.1.40). The following reaction mix was used: 50 mM Hepes pH 7, 10 mM MgCl_2_, 60 mM NH_4_Cl, 250 mM glutamate, 6.25 mM ATP, 1 mM phosphoenolpyruvate, 0.6 mM NADH, 1U PyRK and 1U LDH. Reactions were pre-incubated for 5 minutes at 37 °C and the GS activity was initiated by adding different concentrations of the substrate that was being analyzed. Reactions were developed for 40 min at 37 °C with shaking and absorbance measurement at 340 nm each minute. Analysis of the kinetic characteristics of GS1b.1 WT, GS1b.2 WT and their mutants were performed with GraphPad Prism 8.0.0 (GraphpPad, San Diego, CA, USA).

## ACCESSION NUMBERS

The cDNA sequence data have been submitted to the GenBank database under accession numbers KU641797 (PpGS1a); KU641798, HF548531.1 (PpGS1b.1); and KU641796, KU641799.1, KU641800.1 (PpGS1b.2).

## ACKNOWLEDGEMENTS

The authors are grateful to Professor Francisco Ruiz Cantón for the kind supply of adult tree samples, to Jean François Trontin for the kind supply of embryo samples and to José Miguel Granados for his help during needle harvesting. This work was funded by *Spanish Ministerio de Economía y Competitividad*, grant number BIO2015-73512-JIN MINECO/AEI/FEDER, UE, and *Ministerio de Ciencia e Innovación*, grant number PID2021-125040OB-I00, MICINN, FEDER, UE. This work was also supported by *Junta de Andalucía*, grant number P20_00036 PAIDI 2020/FEDER, UE; and the *Universidad de Málaga* grant B4-2021-01 (Ayudas Plan Propio). JMVM was supported by a scholarship from the Spanish *Ministerio de Educación y Formación Profesional* (FPU17/03517). FO was supported by a grant from the Universidad de Málaga (*Programa Operativo de Empleo Juvenil vía* SNJG, UMAJI11, FEDER, FSE, *Junta de Andalucía*) and funds of the research group BIO-114 (Biología Molecular y Biotecnología) from PAIDI, Junta de Andalucía.

## AUTHOR CONTRIBUTIONS

JMVM, FO and RAC have performed the experiments. RAC performed the phylogenetic analysis. JMVM and JCA have made the *in silico* structural protein analyses. JMVM and RAC have written the manuscript. FO, JCA, CA and FMC made additional contributions and edited the manuscript. JMVM and RAC have planned and designed the research. RAC, CA and FMC were responsible of the funding acquisition.

## DATA AVAILABILITY

Constructions are available from the corresponding authors upon request. The rest of data supporting the findings of this study are available within the paper and within its supplementary materials published online.

## SHORT LEGENDS FOR SUPPORTING INFORMATION

**Dataset S1.** Protein sequences used for phylogenetic analysis.

**Dataset S2.** Phylogenetic distance matrix.

**Dataset S3.** Original tree resulted from phylogenetic analysis in Newick format.

**Table S1.** GSs used for phylogenetic analysis.

**Table S2.** List of primers.

**Figure S1. Promoter region analysis of *PpGS1b.1* and *PpGS1b.2*.** For promoter comparison, 1916 and 2408 nucleotides upstream of start codon of *PpGS1b.1* and *PpGS1b.2* were recovered from genomic data, respectively (Sterck et al., 2022). **A.** Sequences alignment of the promoter region. Position 1995 corresponds to −1 nt before start codon. **B.** Putative transcription factor binding sites identified using the PlantRegMap prediction tool (Tian et al., 2020).

**Figure S2. GS1b.1 WT and GS1b.2 WT quaternary structure. A.** GS1b1.1 WT quaternary structure. **B.** GS1b.2 WT quaternary structure.

**Figure S3. Structural stability of GS1b.1 and GS1b.2 against amino acid substitution.** The presented amino acids have been substituted in each position of GS1b.1 and GS1b.2 amino acid sequences. The differences in the folding free energy between WT and mutant (ΔΔG) for GS1b.1 and GS1b.2 are compared (square plots). The rectangular plots represent the difference between GSb.1 ΔΔG and GS1b.2 ΔΔG (Y axis) for each position (X axis).

**Figure S4. Representation of kinetic characteristics of GS1b.1 WT and GS1b.2 WT for magnesium and ATP. A.** Magnesium. **B.** ATP.

**Figure S5. Subunit structure of the GS1b.1 and GS1b.2 mutants. A.** GS1b.1K264E. **B.** GS1b.1K267H. **C.** GS1b.2E264K. **D.** GS1b.2H267K. Amino acids exchanged, and amino acids associated with them by hydrogen bonds are represented in dark magenta. Amino acids from 330 to the end of the protein are represented in green.

**Figure S6. Quaternary structure of the GS1b.1 and GS1b.2 mutants. A.** GS1b.1K264E. **B.** GS1b.1K267H. **C.** GS1b.1K264E,K267H. **D.** GS1b.2E264K. **E.** GS1b.2H267K. **F.** GS1b.2E264K,H267K.

**Figure S7. Physicochemical properties of the GS1b.1 and GS1b.2 mutants.** Activity of the GS1b.1K264E, GS1b.1K267H, GS1b.2E264K, GS1b.2H267K have been tested at different pH levels (**A**) and temperatures (**B**). Their thermal stability has been also characterized (**C**).

**Figure S8. Representation of kinetic characteristics of GS1b.1 and GS1b.2 single mutants for ammonium.**

**Figure S9. Representation of kinetic characteristics of GS1b.1 and GS1b.2 single and double mutants for glutamate.**

**Figure S10. Representation of kinetic characteristics of GS1b.1 and GS1b.2 single and double mutants for magnesium.**

**Figure S11. Representation of kinetic characteristics of GS1b.1 and GS1b.2 mutants for ATP.**

**Figure S12. Purification of the recombinant GS proteins. A.** Coomassie staining of SDS-PAGE electrophoresis gels. **B.** Western-blot using GS-specific antibodies for GS1b.1 and GS1b.2 detection. Molecular weight ladder (L), soluble fraction (S), binding fraction (B), wash step 1 (W1), wash step 2 (W2), Elution 1 to 9 (E1-9), concentrated fraction (C).

