## Supplementary material for "A new gene encoding a cytosolic glutamine synthetase in pine is linked to developing tissues": Table S2

| Gene | Application | Primer name | Sequence (5’ → 3’) | Primer sense |
| --- | --- | --- | --- | --- |
| *PpGS1a* | Cloning | GS1a-F | TCTCTTGAATCTTATCCCCTTCC | Forward |
|  |  | GS1a-R | TACATATAGCTTATCAGTGACGC | Reverse |
| *PpGS1b.1* | Cloning | GS1b.1-F | CTTCCTCAGGTCGGGCTTG | Forward |
|  |  | GS1b.1-R | CTGAATGACAAACTAGACACTG | Reverse |
| *PpGS1b.2* | Cloning | GS1b.2-F | GCTTCCCCTGCTTTAAGGG | Forward |
|  |  | GS1b.2-R | TGGCACTGACATTCACCAGT | Reverse |
| *PpGS1b.1* | pET30a subcloning | HIS-GS1b.1-F | AGAAATTAATGCACCATCATCATCATCATTCTCTACTGACGGATTTG | Forward |
|  |  | HIS-GS1b.1-R | AGAACTCGAGTCACTTCAAAAGGATGGTTG | Reverse |
| *PpGS1b.2* | pET30a subcloning | HIS-GS1b.2-F | AGAACATATGCACCATCATCATCATCATTCTCTGTTGACAGATTTG | Forward |
|  |  | HIS-GS1b.2-R | AGAACATATGCACCATCATCATCATCATTCTCTGTTGACAGATTTG | Reverse |
| *PpGS1a* | qPCR | qPCR-GS1-F | ATCGAGGAGCTTCAGTTAGAG | Forward |
|  |  | qPCR_GS1a-R | TGGTCGTCTCAGCAATCATAGA | Reverse |
| *PpGS1b.1* | qPCR | qPCR_GS1b.1-F | CCCAATTGTTTGTGGGGGATA | Forward |
|  |  | qPCR_GS1b.1-R | CTGAATGACAAACTAGACACTG | Reverse |
| *PpGS1b.2* | qPCR | qPCR_GS1b.2-F | CCCAATCGTTTCTGTGGATTT | Forward |
|  |  | qPCR_GS1b.2-R | TGGCACTGACATTCACCAGT | Reverse |
| *RNA-binding protein* | qPCR | Pp_27526-F | AAGGCTGTCAAACCTGTCCAA | Forward |
|  |  | Pp_27526-R | TTTAGCTATCAGGATGCCTCTG | Reverse |
| *Saposin-like aspartyl protease* | qPCR | Pp_1135-F | AGTATGCTAAGGAATCGTGCCT | Forward |
|  |  | Pp_1135-R | GTCCATAATTACACACGAACAGA | Reverse |
| *PpGS1b.1* | 264 Mutant | GS1b.1-K264E-F | AAGAAGGGGGAATTGAAGTGATCAAAAAGGCC | Forward |
|  |  | GS1b.1-K264E-R | GGCCTTTTTGATCACTTCAATTCCCCCTTCTT | Reverse |
| *PpGS1b.2* | 264 Mutant | GS1b.2-E264K-F | AAGAAGGTGGAATAAAAGTGATCCATAAGGCC | Forward |
|  |  | GS1b.2-E264K-R | GGCCTTATGGATCACTTTTATTCCACCTTCTT | Reverse |
| *PpGS1b.1* | 267 Mutant | GS1b.1-K267H-F | GGGGGAATTAAAGTGATCCATAAGGCCATT | Forward |
|  |  | GS1b.1-K267H-R | AATGGCCTTATGGATCACTTTAATTCCCCC | Reverse |
| *PpGS1b.2* | 267 Mutant | GS1b.2-H267K-F | GGTGGAATAGAAGTGATCAAGAAGGCCATT | Forward |
|  |  | GS1b.2-H267K-R | AATGGCCTTCTTGATCACTTCTATTCCACC | Reverse |
| *PpGS1b.1* | Double mutant | GS1b.1-264,267-F | GGGGGAATTGAAGTGATCCATAAGGCCATT | Forward |
|  |  | GS1b.1-264,267-R | AATGGCCTTATGGATCACTTCAATTCCCCC | Reverse |
| *PpGS1b.2* | Double mutant | GS1b.2-264,267-F | GGTGGAATAAAAGTGATCAAGAAGGCCATT | Forward |
|  |  | GS1b.2-264,267-R | AATGGCCTTCTTGATCACTTTTATTCCACC | Reverse |

**Table S2.** List of primers.
